# GR chaperone cycle mechanism revealed by cryo-EM: reactivation of GR by the GR:Hsp90:p23 client-maturation complex

**DOI:** 10.1101/2020.09.12.294975

**Authors:** Chari M. Noddings, Ray Yu-Ruei Wang, David A. Agard

## Abstract

Hsp90 is a conserved and essential molecular chaperone responsible for the folding and activation of hundreds of ‘client’ proteins^1,2^. The glucocorticoid receptor (GR) is a model client that constantly depends on Hsp90 for activity^3^. Previously, we revealed GR ligand binding is inhibited by Hsp70 and restored by Hsp90, aided by the cochaperone p23^4^. However, a molecular understanding of the chaperone-induced transformations that occur between the inactive Hsp70:Hsp90 ‘client-loading complex’ and an activated Hsp90:p23 ‘client-maturation complex’ is lacking for GR, or for any client. Here, we present a 2.56Å cryo-EM structure of the GR-maturation complex (GR:Hsp90:p23), revealing that the GR ligand binding domain is, surprisingly, restored to a folded, ligand-bound conformation, while simultaneously threaded through the Hsp90 lumen. Also, unexpectedly, p23 directly stabilizes native GR using a previously uncharacterized C-terminal helix, resulting in enhanced ligand-binding. This is the highest resolution Hsp90 structure to date and the first atomic resolution structure of a client bound to Hsp90 in a native conformation, sharply contrasting with the unfolded kinase:Hsp90 structure^5^. Thus, aided by direct cochaperone:client interactions, Hsp90 dictates client-specific folding outcomes. Together with the GR-loading complex structure (Wang et al. 2020), we present the molecular mechanism of chaperone-mediated GR remodeling, establishing the first complete chaperone cycle for any client.

## Introduction

Hsp90 is required for the functional maturation of 10% of the eukaryotic proteome, including signaling proteins, such as kinases and steroid hormone receptors (SHRs), such as GR^1,6^. We previously uncovered the biochemical basis for GR’s Hsp90 dependence using *in vitro* reconstitution starting with an active GR ligand binding domain (hereafter GR, for simplicity)^4^. We demonstrated that GR ligand binding is regulated by a cycle of GR:chaperone complexes **(Fig. 3d)**. In this chaperone cycle, GR is first inhibited by Hsp70, then loaded onto Hsp90:cochaperone Hop (Hsp70/Hsp90 organizing protein) forming an inactive GR:Hsp90:Hsp70:Hop loading complex (Wang et al. 2020). Upon ATP hydrolysis on Hsp90, Hsp70 and Hop are released, and p23 is incorporated to form an active GR:Hsp90:p23 maturation complex, restoring GR ligand binding with enhanced affinity. Progression through this cycle is coordinated by the ATPase activity of both Hsp70 and Hsp90, which dictate large conformational rearrangements^2,7^. Particularly, Hsp90 functions as a constitutive dimer that undergoes an open-to-closed transition upon ATP binding and this conformational cycle is further regulated by cochaperones^8^. The cochaperone p23 specifically binds and stabilizes the closed Hsp90 conformation^9^ and p23 is required for full reactivation of GR ligand binding *in vitro*^4^ and proper function *in vivo*^10^. Altogether, the coordinated actions of Hsp70, Hsp90, and cochaperones remodel the conformation of GR to control access to the buried, hydrophobic ligand binding pocket.

While the Hsp90/Hsp70 chaperone systems are fundamental in maintaining protein homeostasis, the absence of client:chaperone structures has precluded a mechanistic understanding of the remodeling process for any client. The kinase:Hsp90 structure^5^ first revealed how Hsp90 can stabilize an inactive client, but provided no insights to explain how Hsp90 can reactivate a client, such as GR. Here, we report a high-resolution cryo-EM structure of the GR-maturation complex, providing a long-awaited molecular mechanism for chaperone-mediated client remodeling and activation.

## Results

### Sample preparation and structure determination

The maturation complex sample was prepared through *in vitro* reconstitution of the GR chaperone cycle, whereby MBP (maltose binding protein)-GR ligand binding domain was incubated with Hsp70, Hsp40, Hop, Hsp90, and p23, allowing GR to progress through the chaperone cycle to reach the maturation complex (**Materials and Methods, Supplementary Fig. 1a-d**). A 2.56Å cryo-EM reconstruction of the maturation complex was obtained **(Fig. 1a; Supplementary Fig. 2a,b; Supplementary Table 1)** using RELION^11^ and atomic models were built in Rosetta starting from previously published atomic structures. The structure reveals a fully closed, nucleotide-bound Hsp90 dimer (Hsp90A and B) complexed with a single GR and a single p23, which occupy the same side of Hsp90 **(Fig. 1a,b; Supplementary Fig. 3a)**.

**Figure 1:**
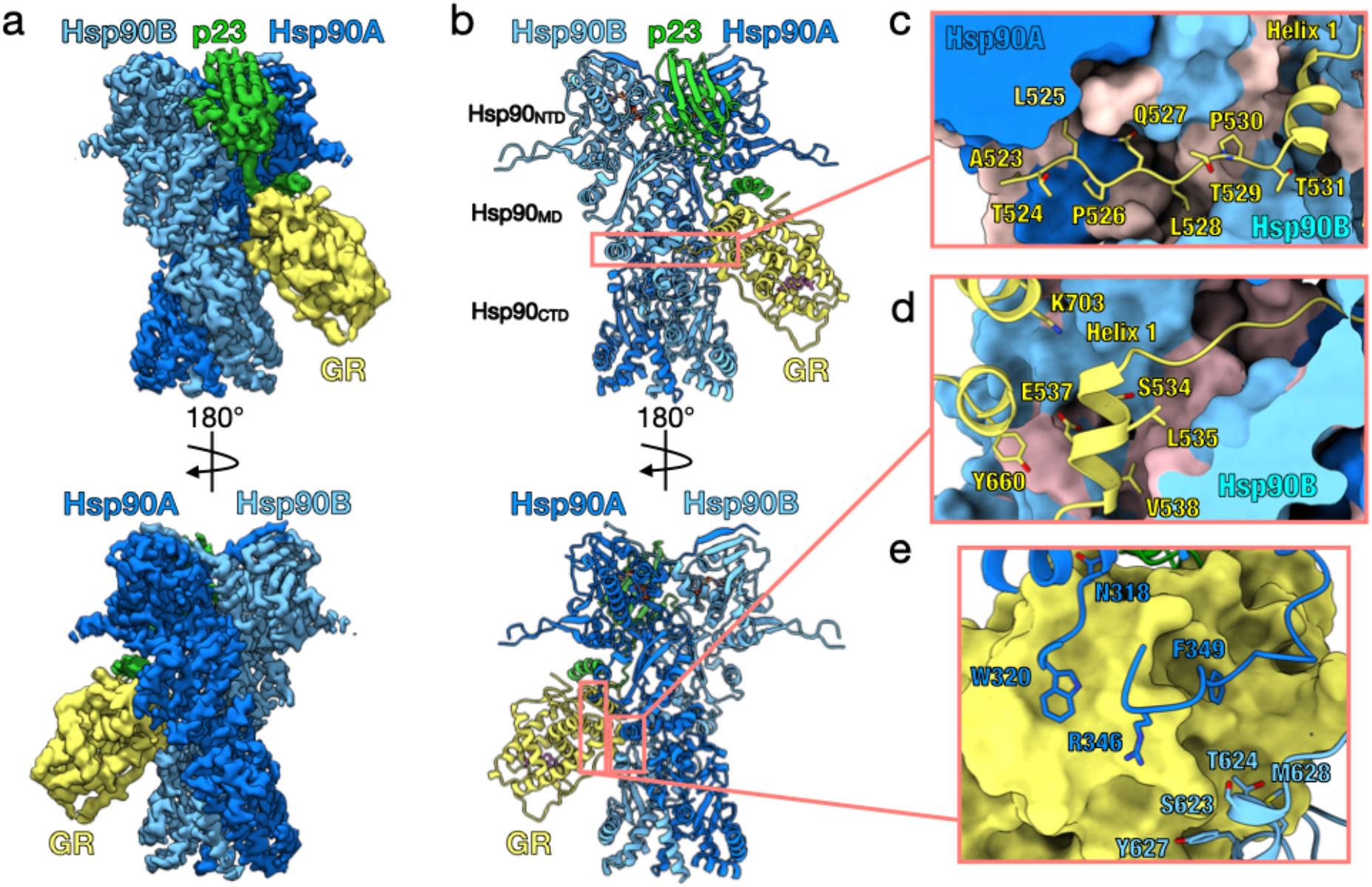
Architecture of the GR-Maturation Complex. **a**, Composite cryo-EM map of the GR-maturation complex. Hsp90A (dark blue), Hsp90B (light blue), GR (yellow), p23 (green). This color scheme is maintained in all figures that show the structure. **b**, Atomic model built into the cryo-EM map. **c**, Interface 1 of the Hsp90:GR interaction depicting the GR_pre-Helix1_ region (GR^523-531^) threading through the Hsp90 lumen. Hsp90A/B are in surface representation. Hydrophobic residues on Hsp90A/B are colored in pink. **d**, Interface 2 of the Hsp90:GR interaction depicting GR_Helix1_ (GR^532-539^) packing against the entrance to the Hsp90 lumen. Hsp90A/B are in surface representation. Hydrophobic residues on Hsp90A/B are colored in pink. **e**, Interface 3 of the Hsp90:GR interaction depicting residues on the Hsp90A_MD_ loops (Hsp90A^N318,W320,R346,F349^) and Hsp90B_amphi-α_ (Hsp90B^S623,T624,Y627,M628^) packing against GR, which is in surface representation.

### Hsp90 stabilizes GR in a native, active conformation

GR and Hsp90 have three major interfaces **(Fig. 1c-e)**: (1) the Hsp90 lumen:GR_pre-Helix1_; (2) the Hsp90_MD/CTD_:GR_Helix1_, and (3) the Hsp90_MD_:GR Helices 3, 4, and 9. In the first interface, the N-terminal GR_pre-Helix1_ (GR^523-531^) is threaded through the closed Hsp90 lumen (∼742Å^2^ buried surface area (BSA))**(Fig. 1c)**. The Hsp90 lumen provides a mostly hydrophobic ‘tunnel’ that captures GR_pre-Helix1_ (**Supplementary Fig. 4a**). Specifically, two residues (GR^L525,L528^) occupy hydrophobic binding pockets within the Hsp90 lumen. The interaction is further stabilized by multiple hydrogen bonds from Hsp90 to the backbone and side chains of GR_pre-Helix1_ **(Supplementary Fig. 4b)**. Interface 2 is mainly comprised of the short GR_Helix1_ (GR^532-539^) packing up against the amphipathic helical hairpin (Hsp90_amphi-α_) in the C-terminal domain of Hsp90B (Hsp90B_CTD_), as well as Hsp90B interactions with GR Helices 8 and 9 (∼425Å^2^ BSA)**(Fig. 1d, Supplementary Fig. 4c)**. In interface 3, the Hsp90B_amphi-α_ packs against GR Helix 3 and the conserved, solvent exposed hydrophobic residues Hsp90^F349,W320^, located at the middle domain of Hsp90A (Hsp90A_MD_), make contact with Helices 4 and 9 on GR (∼421Å^2^ BSA)**(Fig. 1e, Supplementary Fig. 4d)**. Notably, Hsp90^F349,W320^ also make contact with GR in the loading complex (Wang et al. 2020).

Surprisingly, in the maturation complex, despite being bound to Hsp90, GR adopts an active, folded conformation (C*α* RMSD of 1.24Å to crystal structure 1M2Z^12^). Specifically, GR_Helix12_, a dynamic motif responsive to ligand binding, is in the agonist-bound position, as in the crystal structure (1M2Z)**(Supplementary Fig. 5a)**. However, unlike the crystal structure, GR_Helix12_ is not stabilized by a co-activator peptide. Unexpectedly, the density also revealed that GR is ligand-bound **(Supplementary Fig. 5b)**. The only ligand source was the initial GR purification with agonists and GR was extensively dialyzed, which removes the vast majority of ligand^4^. During preparation of the maturation complex, GR likely rebound residual ligand and despite multiple washes, the ligand remained bound, suggesting a slow ligand off-rate from the maturation complex. Based on the ligand density and positions of GR^Y735^, the bound ligand is likely dexamethasone **(Supplementary Fig. 5b)**.

### p23 stabilizes Hsp90 while also interacting directly with GR

The Hsp90:p23 interface is comparable to the crystal structure of the yeast Hsp90:p23 complex (2CG9)^9^, where p23 makes extensive contacts with the N-terminal domains of Hsp90 (Hsp90_NTD_s) to stabilize the Hsp90 closed state (∼1375 Å^2^ BSA**)(Supplementary Fig. 6a-d)**. Only one p23 is bound to the Hsp90 dimer, although two p23 molecules are in the yeast Hsp90:p23 structure and a 2-fold excess of p23 to Hsp90 was added during complex preparation. Consistent with a previous report, GR binding on Hsp90 may favor incorporation of a single p23^13^. The slight asymmetry observed here between the Hsp90_NTD_ dimer interfaces **(Supplementary Fig. 3b)** combined with the avidity afforded by simultaneous interactions with Hsp90 and GR likely provides a molecular explanation.

Unexpectedly, p23 also makes direct and extensive contacts with GR (∼740Å^2^ BSA) through the early part (p23^112-133^) of its ∼57 residue C-terminal tail (p23^104-160^), while the following 27 tail residues (p23^134-160^) were not visible. As seen in the yeast Hsp90:p23 crystal structure (2CG9), the beginning of the p23 tail (p23^F103,N104,W106,D108^) interacts with both Hsp90_NTD_s through multiple hydrogen bonds (**Supplementary Fig. 6c**). The following loop (p23^111-119^) forms multiple hydrogen bonds and salt bridges with both GR and Hsp90 (**Fig. 2a,b**). Although the tail was previously thought to be unstructured^14^, we found an ∼11-residue helix (p23^120-130^, p23_tail-helix_) bound to GR (**Fig. 2a,b; Supplementary Fig. 7a,b**). This newly identified p23_tail-helix_ was also predicted by multiple state-of-the-art secondary structure prediction algorithms (**Supplementary Fig. 7e**). The hydrophobic surface of the p23_tail-helix_ packs against an exposed hydrophobic patch on the GR surface made by helices 9 and 10 **(Supplementary Fig. 7b)**. The p23_tail-helix_ also contacts the C-terminal strand of GR (GR^H775,Q776^), potentially allosterically stabilizing the dynamic GR_Helix12_ **(Fig. 2b, Supplementary Fig. 5c)**.

**Figure 2:**
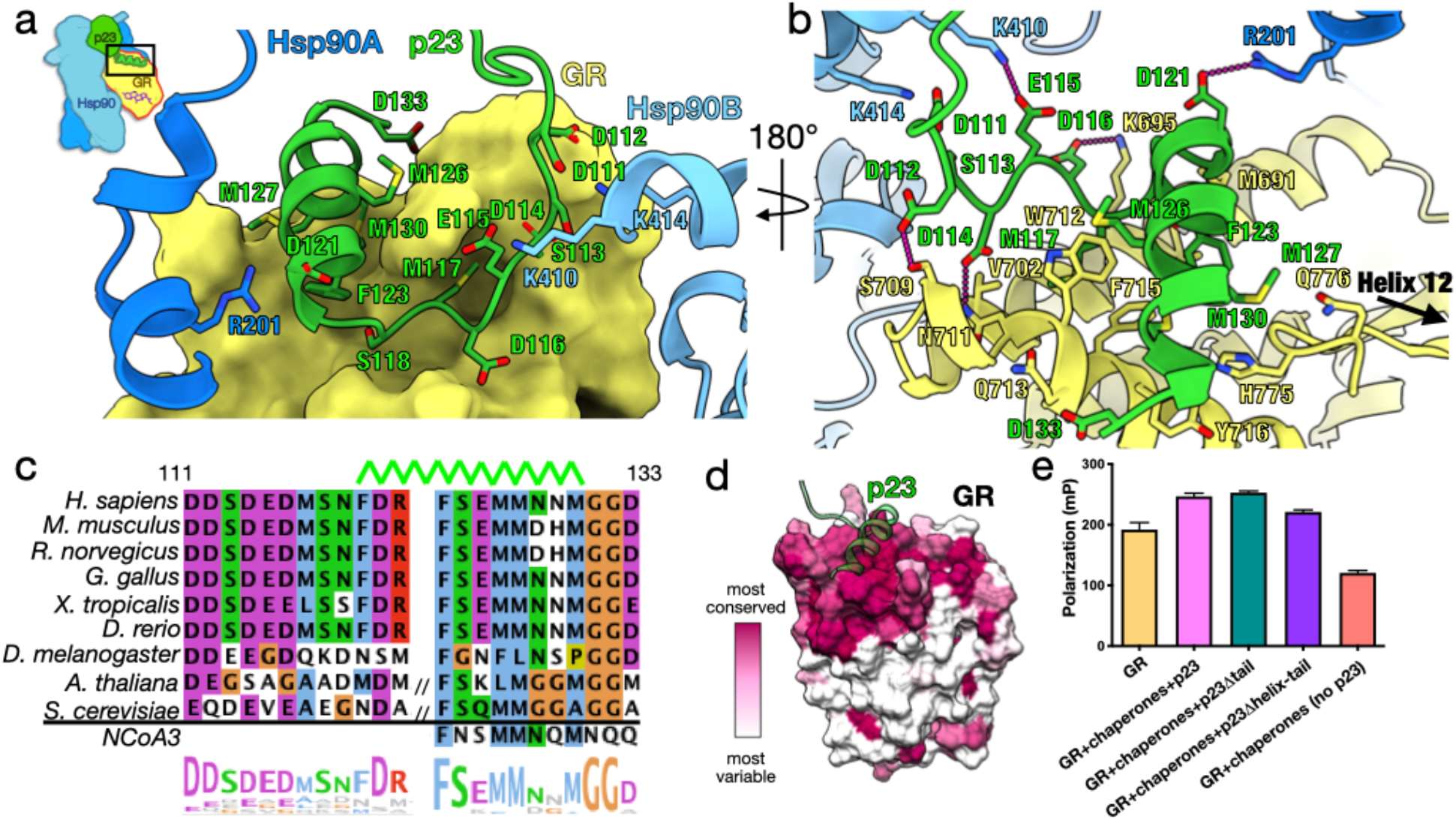
p23_tail-helix_ interactions and effect on GR ligand binding. **a**, Interface between the p23_tail-helix_ (green), GR (yellow, surface representation), and Hsp90 (Hsp90A-dark blue, Hsp90B-light blue). The p23_tail-helix_ (p23^120-130^) binds the top of GR, while the p23 loop (p23^111-119^) interacts with GR and both protomers of Hsp90. **b**, Interface between the p23_tail-helix_ (green), GR (yellow), and Hsp90 (Hsp90A-dark blue, Hsp90B-light blue) showing interacting side chains and hydrogen bonds (dashed pink lines). **c**, Sequence alignment of eukaryotic p23 showing conservation of the p23_tail-helix_ sequence. The top green line indicates the position of the p23_tail-helix_ in the maturation complex atomic model. Slashes indicate insertions in the sequence not shown (longer sequences shown in **Supplementary Fig. 7d**). The alignment is colored according to the ClustalW convention. **d**, Sequence conservation mapped onto GR in surface representation using ConSurf, colored from most variable (white) to most conserved residues (maroon). 87 GR sequences were used for the calculation. The p23_tail-helix_ (light green) was overlaid to indicate the p23:GR interface. **e**, Equilibrium binding of 20nM fluorescent dexamethasone to 250nM GR with chaperone components and p23 tail mutants measured by fluorescence polarization (±SD). Assay conditions were 5 mM ATP, 2 μM Hsp40, 15 μM Hsp70, Hsp90, Hop, and p23/p23 tail mutants.

Notably, in both p23 and GR this novel p23:GR interface is conserved across vertebrates **(Fig. 2c,d)**. The GR hydrophobic patch is also conserved across SHRs, which are thought to undergo similar regulation by Hsp90/Hsp70 **(Supplementary Fig. 7c)**. Attesting to its importance beyond SHRs, the p23_tail-helix_ motif is conserved in yeast, which lack GR; however, there are two additional predicted helices in the much longer yeast p23 tail (**Supplementary Fig. 7d**). Due to the high level of conservation of the p23_tail-helix_ and the hydrophobic patch on SHRs, we reasoned that other proteins may utilize a p23_tail-helix_ -like motif to bind SHRs. Using ScanProsite^15^ to search the human proteome for a p23_tail-helix_ -like motif (“FXXMMN”), remarkably, Nuclear Coactivator 3 (NCoA3/SRC-3), a canonical co-activator protein for SHRs, was among the 10 hits. The identified NCoA3 motif (**F**NS**MM**NQ**M**) aligns with the p23_tail-helix_ sequence and contains the key conserved hydrophobic residues that interact with GR **(Fig. 2c, Supplementary Fig. 7d)**. This suggests NCoA3 may use this newly identified motif to bind GR at the novel interface, in addition to using its LXXLL motif to bind SHRs at Helix 12 (Activation Function 2 (AF-2) Helix)^16^.

### The p23_tail-helix_ is necessary for enhanced GR ligand binding in the chaperone cycle

To quantitatively assess the importance of the p23:GR interface for the enhanced ligand binding in the chaperone cycle, we compared full length p23 to one which lacks both the p23_tail-helix_ and the last 27 C-terminal residues (p23_Δhelix-tail_) and one which only maintains the p23 helix (p23_Δtail_)**(Supplementary Fig. 8a)**. In both cases, the loop between the p23 core and the helix is preserved as this makes critical Hsp90 contacts. While p23_Δtail_ had no significant effect on the chaperone-mediated enhancement of ligand binding, p23_Δhelix-tail_ abolished the enhancement, reducing binding almost to GR alone levels **(Fig. 2e)**. Thus, the observed chaperone-mediated ligand binding enhancement is dependent upon the presence of the p23_tail-helix_. Importantly, p23_Δhelix-tail_ did not reduce the GR ligand binding activity to the same extent as omitting p23, indicating that the p23 core plays a distinct and critical role in stabilizing the closed Hsp90 conformation in the maturation complex. Interestingly, p23 also had an effect on GR ligand binding independent of Hsp90, with addition of p23 to GR modestly increasing ligand binding **(Supplementary Fig. 8b)**.

### Hsp90 contains lumen density in non-GR containing reconstructions

From the same GR:Hsp90:p23 dataset, we also obtained reconstructions of Hsp90:p23 (2.78Å resolution) (**Supplementary Fig. 9a,b)** and MBP:Hsp90:p23 (3.96Å resolution) (**Supplementary Fig. 10a,b)**. In both complexes, the Hsp90 lumen contains density **(Supplementary Fig. 9c, 10d)**. In the MBP:Hsp90:p23 complex, one p23 with low occupancy is bound to Hsp90 on the opposite side of MBP. The MBP is in a partially unfolded state, as density for the two C-terminal helices is missing. The MBP C-terminal region likely threads through Hsp90, accounting for the lumen density. The MBP is also in an apo state, consistent with the unfolding of the last two helices which form part of its binding pocket **(Supplementary Fig. 10c)**.

## Discussion

We present the first atomic resolution structure of a client bound to Hsp90 in a native folded conformation, as well as the highest resolution structure of full-length Hsp90 to date. In the maturation complex, GR simultaneously threads through the closed Hsp90 lumen and adopts a native, ligand-bound conformation that is extensively stabilized by both Hsp90 and the p23_tail- helix_. No GR^apo^ complexes were identified during image analysis, suggesting GR^apo^ is either too dynamic or quickly released from the complex. The native GR conformation in the maturation complex is in striking contrast with the loading complex, in which GR is partially unfolded and unable to bind ligand (Wang et al. 2020). In both complexes, GR threads through the Hsp90 lumen and also interacts on the surface with Hsp90^F349,W320^, although different GR segments are involved. Supporting a general role in client recognition, Hsp90^W320^ is critical for client activation *in vivo*^17-19^. The active, native GR in our complex also starkly contrasts with the only other structure of a closed Hsp90:client complex, which stabilizes an unfolded kinase client^5^. The Hsp90 conformation is nearly identical in both structures and both clients are threaded through the Hsp90 lumen in a similar manner, suggesting a universal binding mode for Hsp90 clients **(Supplementary Fig. 11a,b)**. Although the overall Hsp90:client interactions are similar, the outcomes for folding and function of these two clients are opposing, demonstrating evolutionarily determined, client-specific conformational modulation by Hsp90 **(Supplementary Fig. 11c)**.

While previously thought to be a general cochaperone whose primary function is to stabilize a closed Hsp90, our structure reveals that p23 also makes extensive contacts with GR through a previously uncharacterized helix in the p23 tail. This p23_tail-helix_ is necessary for the observed enhanced GR ligand binding activity *in vitro* and may act by stabilizing ligand-bound GR, securing GR within the complex to indirectly stabilize GR_Helix1_, and/or by allosterically positioning the dynamic GR_Helix12_. Thus, p23 not only serves as a cochaperone to stabilize the closure of Hsp90, but also directly contributes to client maturation. In support of this essential p23:GR interaction, the p23_tail-helix_ and GR hydrophobic groove are well conserved. In fact, the hydrophobic groove is conserved across SHRs, indicating the p23_tail-helix_ may contribute to the Hsp90-dependent chaperoning of all SHRs. Indeed, the activity of all SHRs is dependent on p23^10^ and the progesterone receptor (PR) requires the p23 tail for enhanced ligand binding activity^20^. Intriguingly, NCoA3 contains a p23_tail-helix_ -like motif, suggesting other GR coregulators may utilize this novel helix motif to bind the hydrophobic groove on GR and compete with p23, potentially facilitating GR release. Surprisingly, the p23_tail-helix_ is conserved among eukaryotes that lack SHRs; however, previous studies have demonstrated that the p23 tail has general chaperoning activities^14,20^. To this point, we observed p23 alone could modestly enhance the ligand binding activity of GR independent of Hsp90. Along with the discovery that the Hop cochaperone interacts with the client in the loading complex (Wang et al. 2020), these findings support an emerging paradigm in which Hsp90 cochaperones make specific, direct contact with Hsp90 clients to aid in client recognition and function^5^.

Together with the structure of the GR-loading complex (Wang et al. 2020), we provide for the first time, a complete picture of the chaperone cycle for any client **(Fig. 3d)**. These two structures reveal that GR transitions from a partially unfolded conformation in the loading complex to an active, folded conformation in the maturation complex. In the loading complex, GR_pre-Helix1_ is captured by Hsp70, GR_Helix1_ is stabilized by Hop, and GR_post-Helix1_ is threaded through the semi-closed Hsp90 lumen. First, Hsp70 releases and then Hop releases, which lets GR_pre-Helix1_ slide into the Hsp90 lumen, allowing GR_Helix1_ to refold onto the GR core, thereby generating a ligand binding capable, native GR, stabilized by the p23_tail-helix_ **(Fig. 3b)**. During this transition, Hsp90 twists to the fully closed conformation, which likely facilitates client sliding and helps rearrange the client binding site to fully enclose GR_pre-Helix1_ **(Fig. 3a)**. Perhaps most critical for GR function, GR becomes protected from Hsp70 rebinding and inhibition once it is in the maturation complex. GR_Helix1_ has been proposed to function as a lid over the GR ligand binding pocket^21^, thus ligand likely binds during the transition from the loading complex to the maturation complex, just as GR_Helix1_ slides through the Hsp90 lumen to seal the ligand binding pocket **(Fig. 3c)**. In line with previous studies, our findings suggest that ligand-bound GR may be translocated to the nucleus in the maturation complex^22^, perhaps with the aid of FKBP52^23^, where it would be protected by Hsp90 from re-inhibition by Hsp70. Supporting this, Hsp90 and p23 have been found in the nucleus colocalized with GR^24^, suggesting the maturation complex may persist even after translocation.

**Figure 3:**
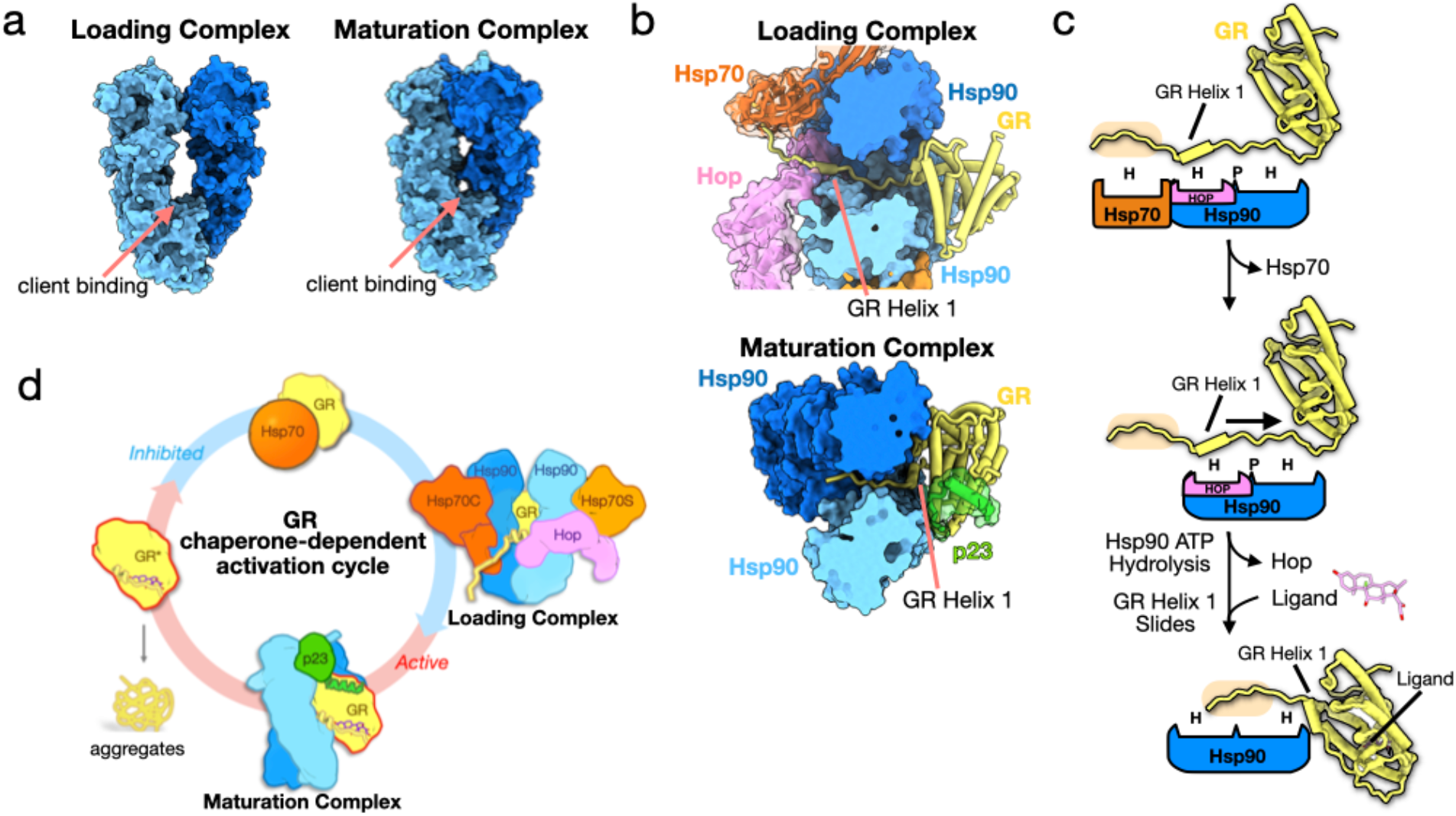
Mechanism of GR Activation by Hsp90. **a**, Surface representation of Hsp90 in the GR-loading complex (Wang et al. 2020) versus the GR-maturation complex showing the change in Hsp90 conformation. Hsp90A (dark blue), Hsp90B (light blue) **b**, Top view of the loading complex (top), where GR_Helix1_ is extended through the Hsp90 lumen, versus a top view of the maturation complex (bottom), where GR_Helix1_ is docked back onto GR. Hsp70 (orange, partially transparent surface), Hop (pink, partially transparent surface), Hsp90A (dark blue, surface representation), Hsp90B (light blue, surface representation), GR (yellow), p23 (green, partially transparent surface). **c**, Schematic showing the conformational change of the GR_Helix1_ region from the loading complex to the maturation complex. Boxes represent chaperone and co-chaperone binding sites along the GR_Helix1_ region (H=hydrophobic interface, P=polar interface). The GR_pre-Helix1_ strand is highlighted (tan). Color scheme is maintained from (**b**). **d**, Schematic of the GR chaperone-dependent activation cycle. Ligand-bound, active GR (left) is aggregation prone under physiological conditions. GR ligand binding is inhibited upon binding to Hsp70. Inactive GR is then loaded onto Hsp90, with Hop, to form the loading complex, where GR_Helix1_ is extended through the Hsp90 lumen. Hsp70 and Hop leave and p23 is incorporated to form the maturation complex, where GR_Helix1_ slides back onto the body of GR and ligand binding is restored.

The proposed sliding mechanism may be a general theme for Hsp90’s client remodeling mechanism. Our two other reconstructions, Hsp90:p23 and MBP:Hsp90:p23, have density in the Hsp90 lumen, suggesting Hsp90 has bound regions in our construct other than GR. Given that GR_Helix1_ slides through the Hsp90 lumen from the loading complex to the maturation complex, it is possible that Hsp90 can act processively during open-to-closed transitions to remodel other client domains beyond the one initially engaged. This would explain these other structural classes, although this remains to be tested. Nevertheless, the mechanism of GR_Helix1_ sliding explains how Hsp90 can provide protected refolding of client domains as they exit the lumen to become directly stabilized by cochaperones. Our results suggest Hsp90 may use this mechanism to allow domains to fold independently on either side of the lumen or uncouple annealed misfolded regions to ensure folding fidelity.

## Acknowledgements

We thank members of the Agard Lab and Elaine Kirschke for helpful discussions. We thank David Bulkley, Glenn Gilbert, and Zanlin Yu from the W.M. Keck Foundation Advanced Microscopy Laboratory at the University of California, San Francisco (UCSF) for EM facility maintenance and help with data collection. We also thank Matt Harrington and Joshua Baker-LePain for computational support with the UCSF Wynton cluster. R.Y.-R.W is a Howard Hughes Medical Institute Fellow of the Life Sciences Research Foundation. This work was supported by funding from Howard Hughes Medical Institute and National Institutes of Health grants R35GM118099, S10OD020054, S10OD021741 (to D.A.A.).

## Author Contributions

C.M.N. and R.Y.-R.W. designed and executed biochemical experiments, cryo-EM sample preparation, data collection, data processing, and model building. C.M.N., R.Y.-R.W., and D.A.A. conceived the project, interpreted the results, and wrote the manuscript.

