## Supplementary Materials for "GR chaperone cycle mechanism revealed by cryo-EM: reactivation of GR by the GR:Hsp90:p23 client-maturation complex"

This PDF file includes:

Supplementary Figures 1 to 11; Supplementary Table 1

Materials and Methods

References for Materials and Methods

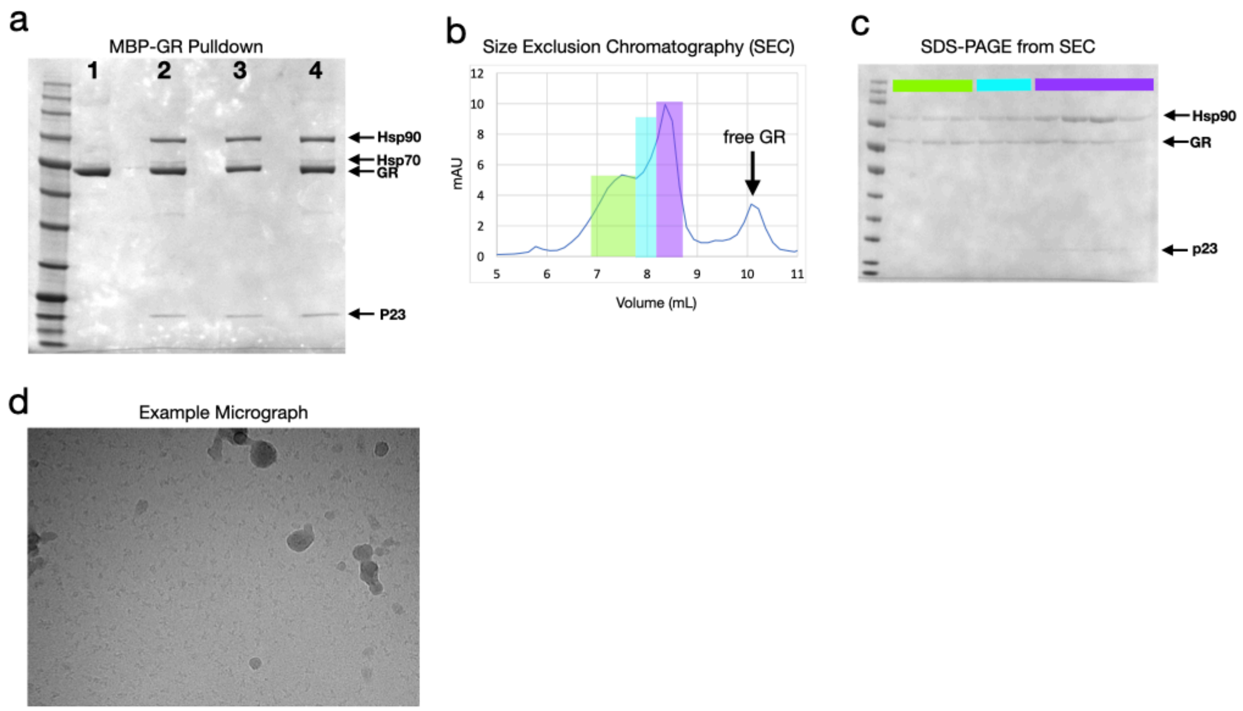


**Supplementary Figure 1: Sample Preparation**

**a,** Coomassie-stained SDS-PAGE with elution from the MBP-GR pulldown from the *in vitro* reconstituted GR chaperone cycle. Assay conditions are as follows- Lane 1:5uM MBP-GR; Lanes 2-4: 5uM MBP-GR, 2uM Hsp40, 5uM Hsp70, 5uM Hop, 15uM Hsp90, 15uM Bag-1, 30uM p23, 5mM ATP, 20mM molybdate. **b,** Shodex KW-804 gel-filtration profile of the GR maturation complex purified by MBP-GR pulldown from the reconstituted GR chaperone cycle. **c,** Coomassie-stained SDS-PAGE of the fractions from gel filtration. Colors indicate which gel lanes correspond to specific regions of the gel-filtration profile. Sample fractions from the region highlighted in purple were collected and used for cryo-EM analysis. **d,** Representative electron micrograph for the cryo-EM dataset.

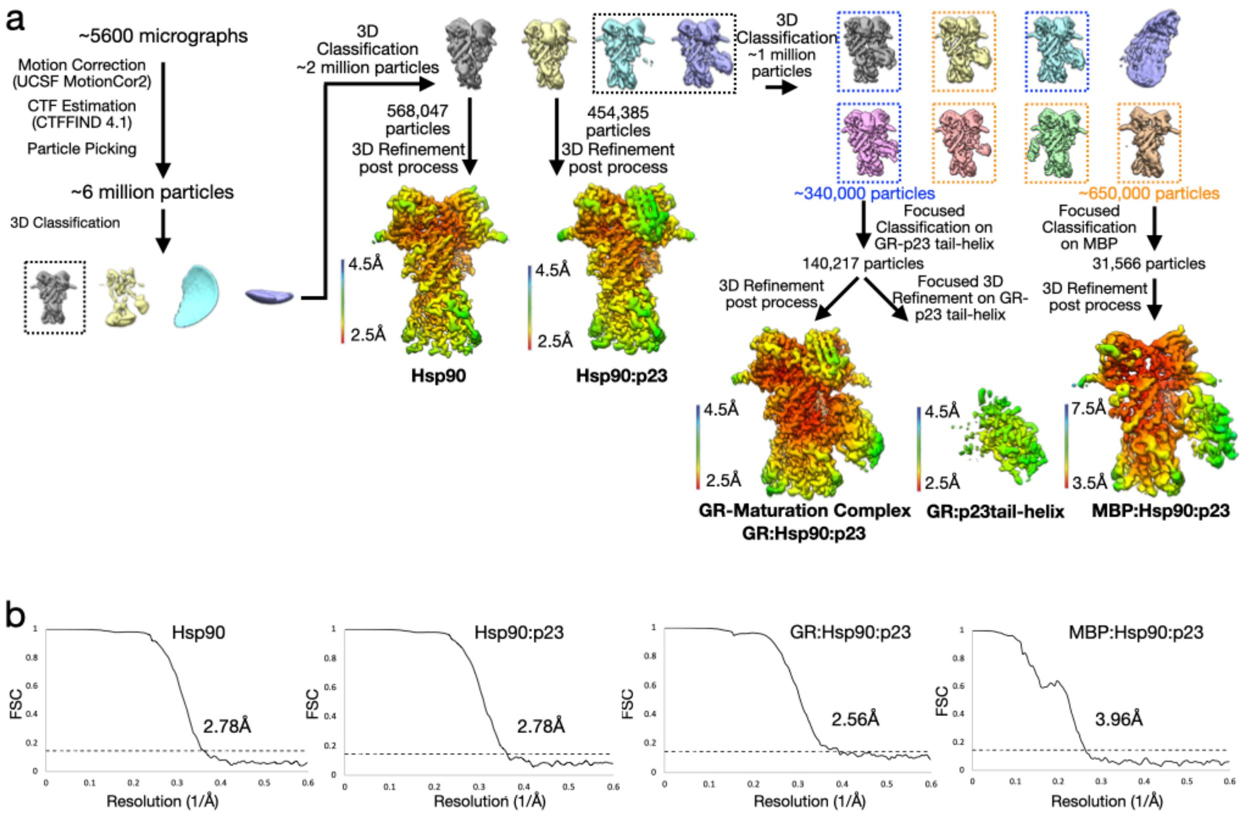


**Supplementary Figure 2: Cryo-EM Data Analysis**

**a,** Cryo-EM data processing procedure with 3D maps colored by local resolution. **b,** Gold-standard Fourier shell correlation (FSC) curves of the 3D reconstructions. The dashed lines intercept the y axis at an FSC value of 0.143.

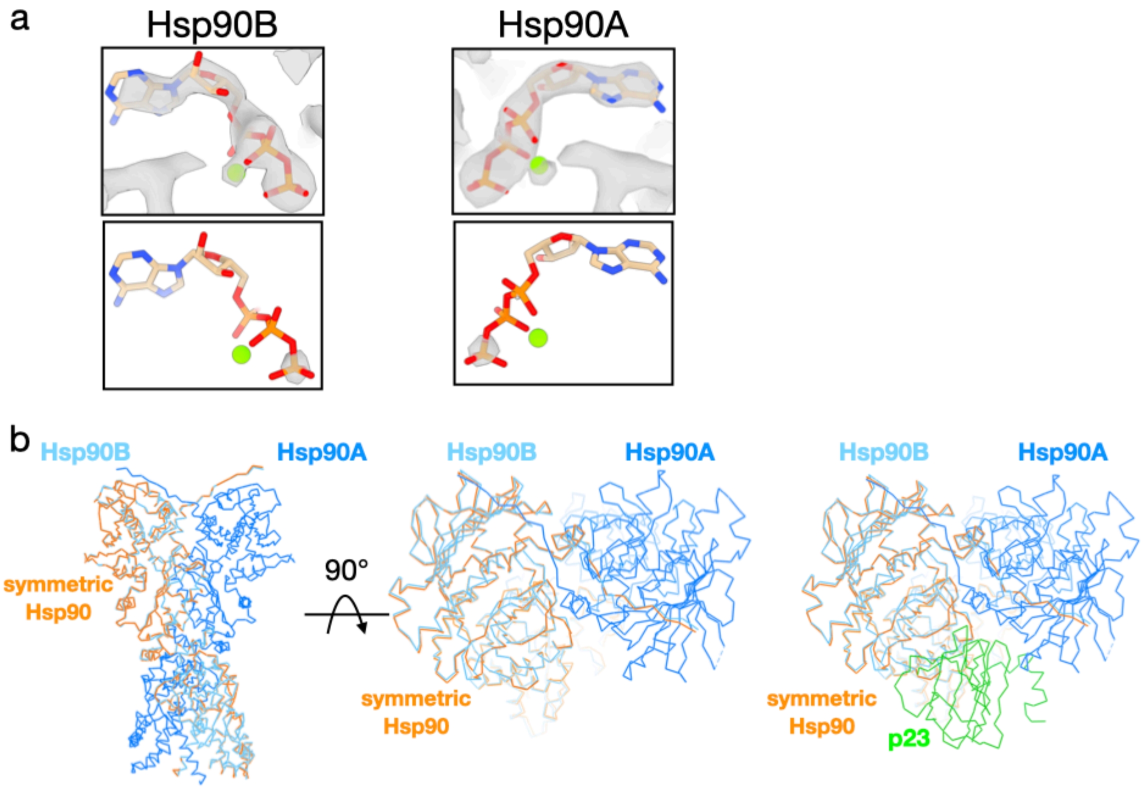


**Supplementary Figure 3: Hsp90 Conformation and Nucleotide-Bound State**
**a,** GR maturation complex map density with atomic model showing ATP-magnesium density in both Hsp90 protomers (Hsp90A/B). Bottom images show increased threshold on the map density to indicate that the ATP position has relatively strong density, likely corresponding to molybdate (see Methods). **b,** Atomic model of a symmetric Hsp90 dimer (orange) compared with Hsp90 from the maturation complex atomic model, indicating a slight asymmetry in the Hsp90 dimer interface in the maturation complex. Hsp90A (dark blue), Hsp90B (light blue), p23 (green).

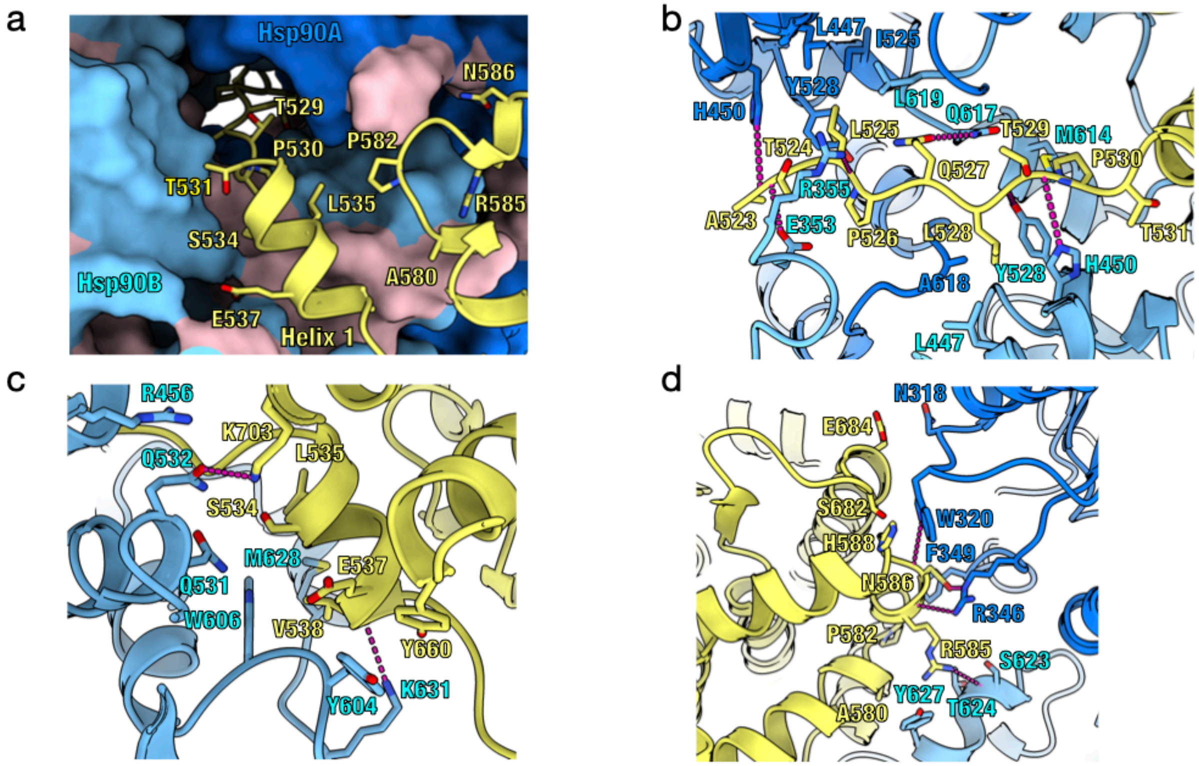


**Supplementary Figure 4: Hsp90-GR Interfaces**

Atomic model of the maturation complex with Hsp90A (dark blue), Hsp90B (light blue), GR (yellow). **a,** View of the $GR_{pre\text{-}helix\ 1}$ strand threaded through the Hsp90 lumen and GR helices 1 and 3 packing against the entrance to the Hsp90 lumen. Side chains on GR in contact with Hsp90 are shown. Hsp90A/B are in surface representation. Hydrophobic residues on Hsp90 are colored in pink. **b,** Interface 1 of the Hsp90:GR interaction depicting the $GR_{pre\text{-}Helix\ 1}$ region ($GR^{523\text{-}531}$) threading through the Hsp90 lumen. Side chains in contact between GR and Hsp90 are shown, along with hydrogen bonds (dashed pink lines). **c,** Interface 2 of the Hsp90:GR interaction depicting $GR_{Helix\ 1}$ ($GR^{532\text{-}539}$) packing against Hsp90. Side chains in contact between GR and Hsp90 are shown, along with hydrogen bonds (dashed pink lines). **d,** Interface 3 of the Hsp90:GR interaction depicting residues on the $Hsp90A_{MD}$ loops ($Hsp90A^{N318,W320,F349,R346}$) and $Hsp90B_{amphi\text{-}\alpha}$ ($Hsp90B^{S623,T624,Y627,M628}$) packing against GR. Side chains in contact between GR and Hsp90 are shown, along with hydrogen bonds (dashed pink lines).

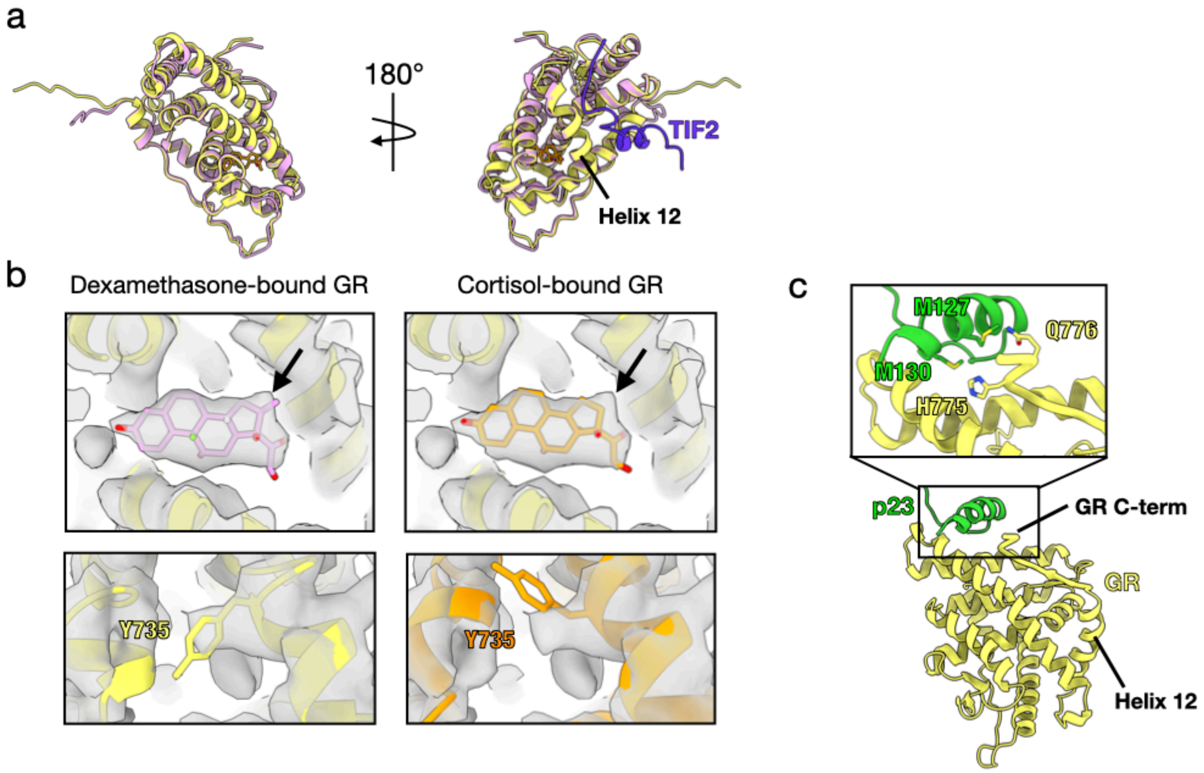


**Supplementary Figure 5: GR is in a Native, Ligand-Bound Conformation**
**a,** Atomic model of GR from the maturation complex (yellow) compared with GR from the crystal structure (1M2Z; GR, pink; co-activator peptide TIF2, purple; ligand, orange). $GR_{Helix\ 12}$ is indicated. **b,** GR maturation complex map density with atomic models. In the top images, the ligand density is shown with either the agonist dexamethasone docked (left) or the agonist cortisol docked (right). Arrow indicates the extra carbon atom in dexamethasone compared to cortisol. In the bottom images, density for $GR^{Y735}$ is shown with either the dexamethasone-bound crystal structure docked (left image, 1M2Z) or the cortisol-bound crystal structure docked (right image, 4P6X). **c,** Atomic model of GR (yellow) and p23 (green) from the maturation complex highlighting the interaction between the $p23_{tail\text{-}helix}$ and the GR C-terminus, which connects to $GR_{Helix\ 12}$.

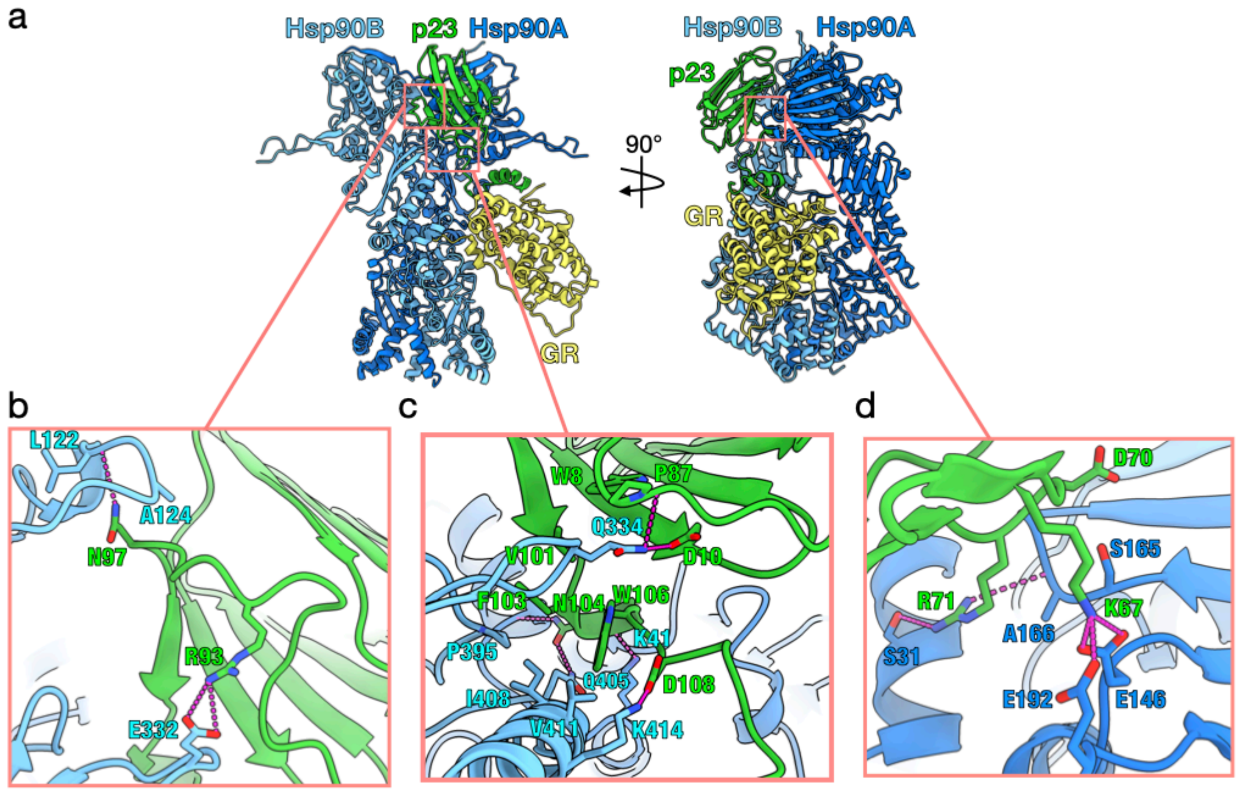


**Supplementary Figure 6: Hsp90-p23 Interfaces**

Atomic model of the maturation complex with Hsp90A (dark blue), Hsp90B (light blue), GR (yellow), p23 (green). **b,** Interface 1 of the Hsp90:p23 interaction depicting Hsp90B interacting with one side of the p23 core. Side chains in contact between p23 and Hsp90B are shown, along with hydrogen bonds (dashed pink lines). **c,** Interface 2 of the Hsp90:p23 interaction depicting Hsp90A and Hsp90B interacting with the base of the p23 core. Side chains in contact between p23 and Hsp90A/B are shown, along with hydrogen bonds (dashed pink lines). **d,** Interface 3 of the Hsp90:p23 interaction depicting Hsp90A interacting with the side of the p23 core. Side chains in contact between p23 and Hsp90 are shown, along with hydrogen bonds (dashed pink lines).

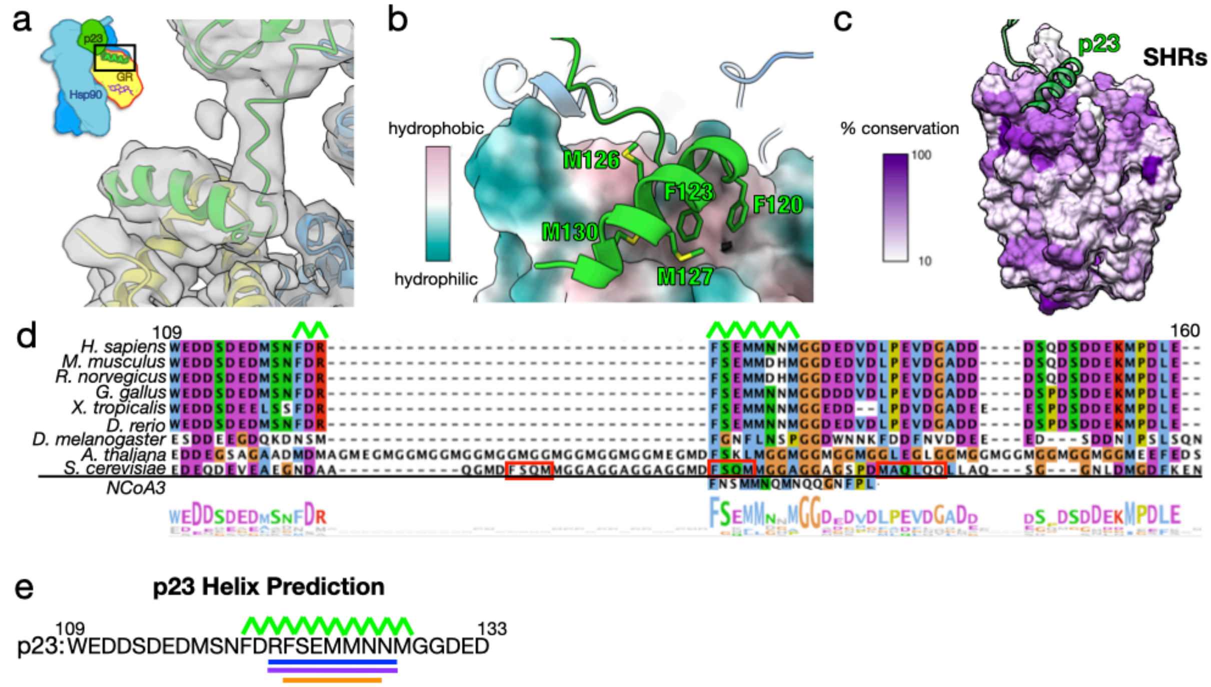


**Supplementary Figure 7: The p23$_{\text{tail-helix}}$-GR interface**

**a,** Cryo-EM map showing density for the p23 tail with the atomic model built in. Hsp90A (dark blue), Hsp90B (light blue), GR (yellow), p23 (green). **b,** Interface between the p23$_{\text{tail-helix}}$ (green) and GR (colored by hydrophobicity, surface representation) showing the p23$_{\text{tail-helix}}$ binds to a hydrophobic patch on GR. p23 side chains interacting with GR are shown. **c,** Sequence identity across human steroid hormone receptors (GR, mineralocorticoid receptor, androgen receptor, progesterone receptor, estrogen receptor α and β) plotted onto the GR structure. The p23$_{\text{tail-helix}}$ (light green) was overlaid to indicate the p23:GR interface. **d,** Sequence alignment of eukaryotic p23 showing conservation of the p23$_{\text{tail-helix}}$ sequence. The bottom aligned sequence is the p23$_{\text{tail-helix}}$ -like motif identified in NCoA3 using the ScanProsite server. Red boxes on the *S. cerevisiae* p23 sequence indicate predicted helices from the PsiPred server. The alignment is colored according to the ClustalW convention. **e,** Secondary structure predictions for human p23 from three different servers. Psipred (purple), Porter 4.0 (orange), RaptorX (blue). The p23$_{\text{tail-helix}}$ from the maturation complex atomic model is shown with the top green lines.

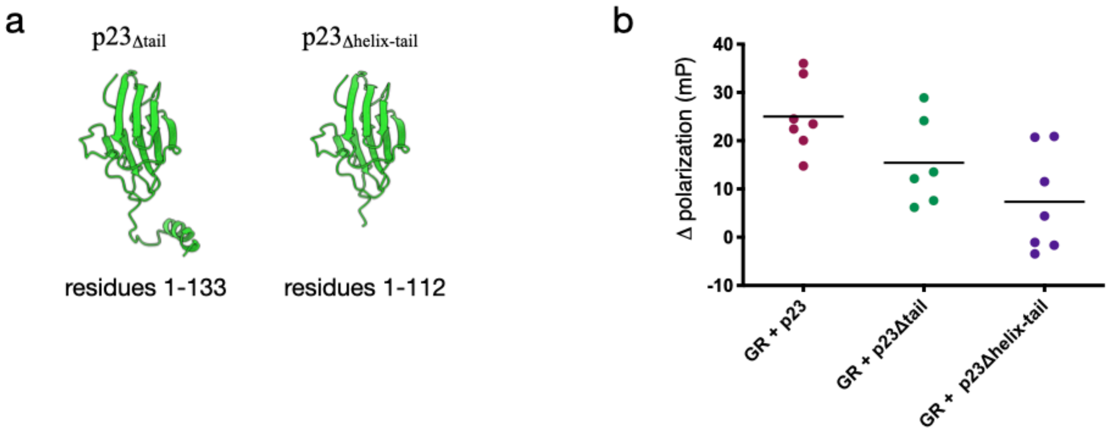


**Supplementary Figure 8: Effect of p23 Tail Mutants on GR Ligand Binding**
**a,** Depiction of the two p23 tail mutants ($p23_{\Delta tail}$, residues 1-133; $p23_{\Delta helix\text{-}tail}$, residues 1-112) used in the GR ligand binding assays. **b,** Equilibrium binding of 20nM fluorescent dexamethasone to 250nM GR with addition of 15uM p23 or p23 tail mutants measured by fluorescence polarization (±SD). Polarization values were normalized to the equilibrium binding of 20nM fluorescent dexamethasone to 250nM GR without p23/p23 tail mutants.

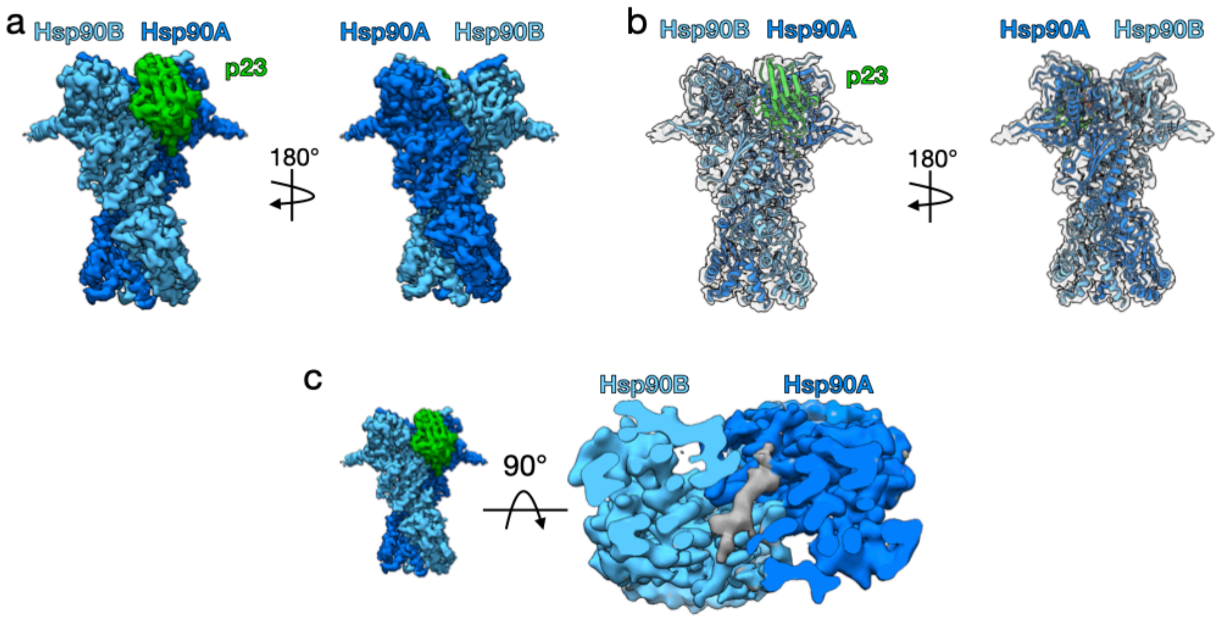


**Supplementary Figure 9: Hsp90-p23 Complex**
**a,** Cryo-EM map of the Hsp90:p23 complex. Hsp90A (dark blue), Hsp90B (light blue), p23 (green). This color scheme is maintained in all figures that show the structure. **b,** Atomic model of Hsp90 and p23 from the GR-maturation complex docked into the Hsp90:p23 map density. **c,** Top view of the Hsp90:p23 complex density map with clipping plane to show unidentified density (gray) through the Hsp90 lumen.

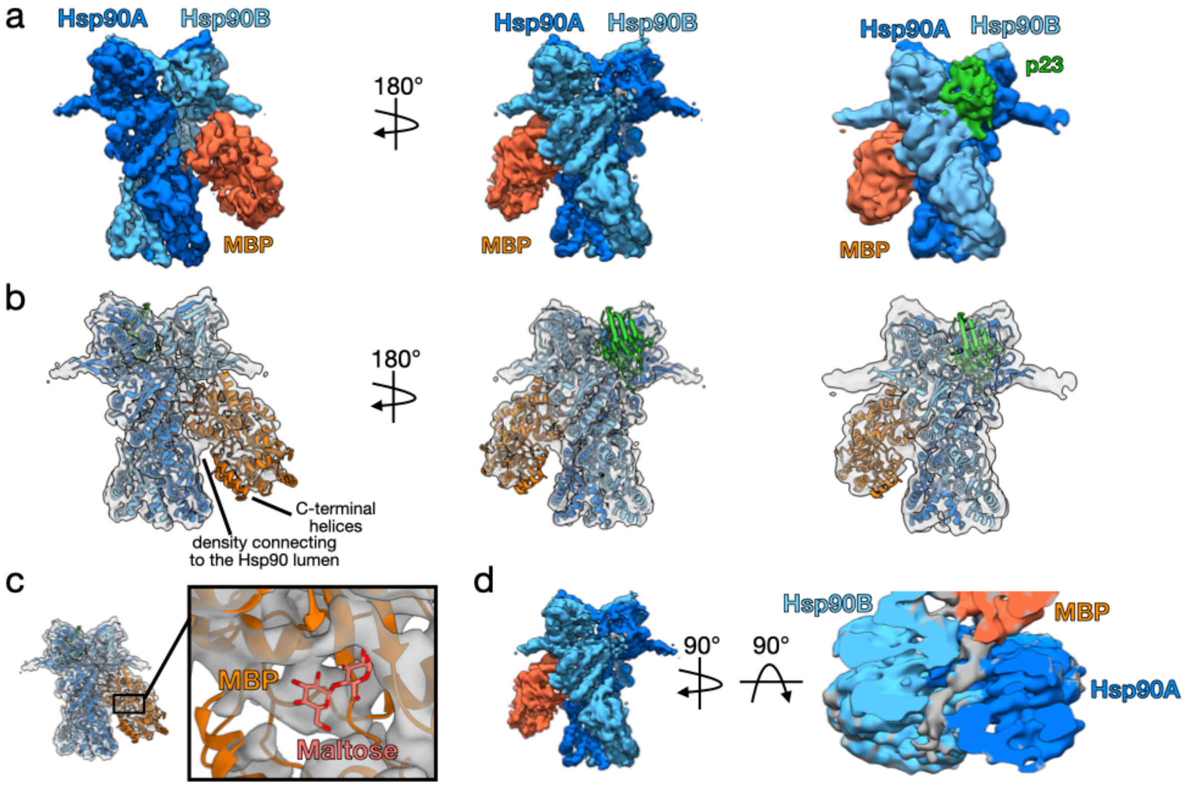


**Supplementary Figure 10: MBP-Hsp90-p23 Complex**
**a,** Cryo-EM map of the MBP:Hsp90:p23 complex. Far right image shows the lowpass-filtered density map. Hsp90A (dark blue), Hsp90B (light blue), p23 (green), MBP (orange). This color scheme is maintained in all figures that show the structure. **b,** The apo MBP crystal structure (1OMP) and atomic model of Hsp90 and p23 from the GR-maturation complex are docked into the MBP:Hsp90:p23 map density. Far right image shows the density map lowpass-filtered to 6Å. **c,** Maltose-bound MBP crystal structure (1ANF) docked into the MBP:Hsp90:p23 map density. MBP (orange), maltose (pink). **d,** Top view of the MBP:Hsp90:p23 complex density map with clipping plane to show unidentified density (gray) through the Hsp90 lumen.

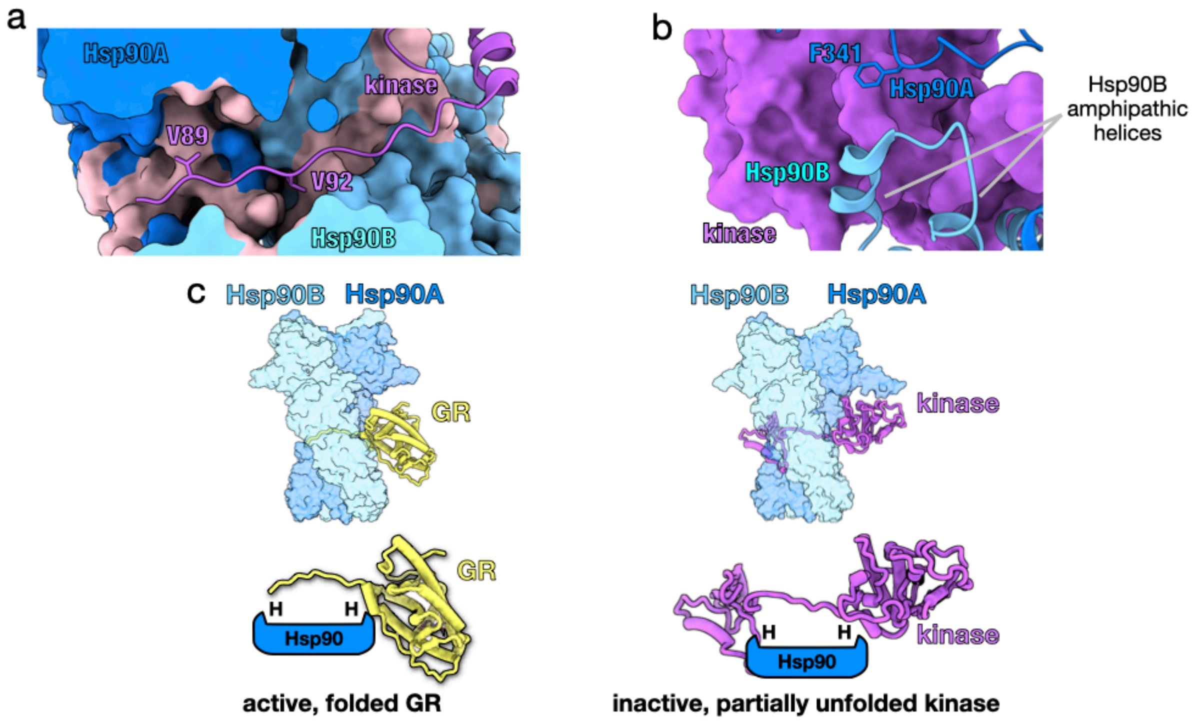


**Supplementary Figure 11: Comparison of the GR-Maturation Complex with the Hsp90-Kinase Complex**

**a,** Structure of Hsp90 bound to an unfolded kinase client (5FWK) depicting a strand of the kinase client threaded through the Hsp90 lumen. The two hydrophobic residues on the kinase ($Cdk4^{V89,V92}$) that occupy the Hsp90 hydrophobic pockets in a similar manner to the GR-maturation complex are displayed. Hsp90A (dark blue), Hsp90B (light blue), Cdk4 kinase (purple). Hsp90A/B are in surface representation. Hydrophobic residues on Hsp90 are colored in pink. **b,** Structure of Hsp90 bound to an unfolded kinase client (5FWK) depicting $Hsp90A^{F341}$ ($Hsp90A^{F349}$ in the GR-maturation complex) and $Hsp90B_{amphi\text{-}\alpha}$ packing against the kinase. Hsp90A (dark blue), Hsp90B (light blue), Cdk4 kinase (purple, surface representation). **c,** Top images are structures of the GR-maturation complex and Hsp90:kinase complex showing both clients thread through the closed Hsp90 lumen. Hsp90A (dark blue), Hsp90B (light blue), GR (yellow), Cdk4 kinase (purple). Hsp90A/B are in surface representation. Bottom images are schematics demonstrating that both clients thread through the mostly hydrophobic Hsp90 lumen, but display different folding outcomes (H=hydrophobic interface).

| | GR:Hsp90:p23 | Hsp90 | Hsp90:p23 | MBP:Hsp90:p23 |
|---|---|---|---|---|
| **Data Collection and Processing** | | | | |
| Magnification | 105,000 | 105,000 | 105,000 | 105,000 |
| Voltage (kV) | 300 | 300 | 300 | 300 |
| Electron Exposure ($e^-/Å^2$) | 60 | 60 | 60 | 60 |
| Defocus range (μm) | 0.8-2.0 | 0.8-2.0 | 0.8-2.0 | 0.8-2.0 |
| Pixel size (Å) | 0.835 | 0.835 | 0.835 | 0.835 |
| Symmetry imposed | C1 | C1 | C1 | C1 |
| Initial particles (no.) | ~6,000,000 | ~6,000,000 | ~6,000,000 | ~6,000,000 |
| Final particles (no.) | 140,217 | 568,047 | 454,385 | 31,566 |
| Map resolution (Å) | 2.56 | 2.78 | 2.78 | 3.96 |
| FSC threshold | 0.143 | 0.143 | 0.143 | 0.143 |
| Map resolution range (Å) | 2.5-4.5 | 2.5-4.5 | 2.5-4.5 | 3.5-5.5 |

**Supplementary Table 1: Cryo-EM Data Collection**

### Materials and Methods

#### *Data analysis and figure preparation*

Figures were created using UCSF Chimera v.1.14[1] and UCSF ChimeraX v0.94[2]. GR ligand binding data was analyzed using Prism v.8.4.3 (GraphPad).

#### *Protein expression and purification*

Human Hsp90α, Hsp70 (Hsp70A1A), Hop, p23, p23$_{\Delta tail}$ (1-133), p23$_{\Delta helix\text{-}tail}$ (1-112), and yeast Ydj1 were expressed in the pET151 bacterial expression plasmid with a cleavable N-terminal, 6x-His tag. Human Bag-1 isoform 4 (116-345) was expressed in a pET28a vector with a cleavable N-terminal, 6x-His tag. Proteins were expressed and purified by the following procedure. Proteins were expressed in bacterial BL21 star (DE3) strain. Cells were grown in either LB or TB at 37°C until $OD_{600}$ reached 0.6-0.8 and then induced with 0.5 mM IPTG overnight at 16°C. Cells were harvested and lysed in 50 mM Potassium Phosphate pH 8, 500 mM KCl, 10 mM imidazole pH 8, 10% glycerol, 6 mM βME, and Roche cOmplete, mini protease inhibitor cocktail using an EmulsiFlex-C3 (Avestin). Lysate was centrifuged and the soluble fraction was affinity purified by gravity column with Ni-NTA affinity resin (QIAGEN). The protein was eluted with 30 mM Tris pH8, 50 mM KCl, 250 mM imidazole pH 8, and 6 mM βME. For Hsp90, Hsp70, and Ydj1, an extra wash step with 0.1% Tween20 and 2 mM ATP/$MgCl_2$ was added to the Ni-NTA resin before eluting. The 6x-His tag was removed with TEV protease during the following overnight dialysis in 30 mM Tris pH 8, 50 mM KCl, and 6 mM βME. Cleaved protein was then loaded onto an ion exchange column, MonoQ 10/100 GL (GE Healthcare), with 30 mM Tris pH 8, 50 mM KCl, and 6 mM βME and eluted with a linear gradient of 50-500 mM KCl. Protein was further purified by size exclusion in 30 mM HEPES pH

7.5, 50 mM KCl, 10% glycerol, 1 mM DTT using a HiLoad 16/60 Superdex 200 (GE Healthcare) or Hi Load 16/60 Superdex 75 (GE Healthcare). For Hsp70, each peak from ion exchange was collected separately and purified by size exclusion in 30 mM HEPES pH 7.5, 100 mM KCl, 10% glycerol, 4 mM DTT, where only the monomeric peak was then collected. Protein was concentrated, flash frozen, and stored at -80$^{\circ C}$.

*GR-LBD expression and purification*

For GR-LBD, LBD(F602S)(521-777) was codon optimized and expressed in the pMAL-c3X derivative with an N-terminal cleavable 6x-His-MBP tag. GR-LBD was expressed and purified as previously described[3].

*GR-maturation complex sample preparation*

The GR chaperone cycle was reconstituted *in vitro* with purified components as previously described[3]. Buffer conditions were 30 mM HEPES pH 8, 50 mM KCl, 0.05% Tween20, and 2 mM TCEP. Proteins and reagents were added at the following concentration: 5 μM MBP-GR LBD, 2 μM Hsp40, 5 μM Hsp70, 5 μM Hop, 15 μM Hsp90, 15 μM p23, 5 mM ATP/$MgCl_2$. This reaction was incubated at room temperature for 60 minutes, then 15 μM p23, 15 μM Bag-1, and 20 mM sodium molybdate (used to stabilize the closed conformation of Hsp90[4,5]) were added, and the reaction was incubated at room temperature for another 30 minutes. Following incubation, amylose resin (New England Biolabs) was added the reactions in a 1:1 ratio and incubated at 4°C with nutation. Resin was then washed 4 times with wash buffer (30 mM HEPES pH 8, 50 mM KCl, 5 mM ATP/$MgCl_2$, 0.05% Tween20, 2 mM TCEP, 20 mM sodium molybdate) and eluted with 50 mM maltose. The elution was analyzed by SDS-PAGE

(**Supplementary Fig.1a**). The elution was concentrated and purified by size exclusion using a Shodex KW-804 on an Ettan LC (GE Healthcare)(**Supplementary Fig.1b,c**). Fractions containing the full complex were concentrated to ~2 μM. 2.5 μL of sample was applied to glow-discharged QUANTIFOIL R1.2/1.3, 400-mesh, copper holey carbon grid (Quantifoil Micro Tools GmbH) and plunge-frozen in liquid ethane using a Vitrobot Mark IV (FEI) with a blotting time of 15 seconds, at 10°C, and with 100% humidity.

*Cryo-EM data acquisition*

The images were collected on a FEI Titan Krios electron microscope (Thermo Fisher Scientific) operating at 300kV using a K3 direct electron camera (Gatan) (example micrograph **Supplementary Fig. 1d**). Images were recorded at a nominal magnification of 105,000×, corresponding to a physical pixel size of 0.835Å. A nominal defocus range of 0.8 μm –2.0 μm underfocus was used. A total exposure of 5.9 seconds was used with 0.05 second subframes (117 total frames). The total accumulated electron dose was 60 electrons/$Å^2$ and 0.5128 electrons/$Å^2$/frame. Data was acquired using SerialEM software[6].

A small dataset on the GR-maturation complex was collected before the larger dataset described above. The smaller dataset was collected from the same GR-maturation complex sample preparation concentrated to 1.2 μM with grids prepared in a similar manner. Images were collected on a FEI Titan Krios electron microscope (Thermo Fisher Scientific) operating at 300kV using a K3 direct electron camera (Gatan). Images were recorded at a nominal magnification of 105,000×, corresponding to a physical pixel size of 0.835Å. A nominal defocus range of 0.8 μm –2.0 μm underfocus was used. A total exposure of 3.0 seconds was used with 0.0255 second subframes (118 total frames). Data was acquired using SerialEM software.

*Cryo-EM data processing*

The smaller dataset consisted of ~1500 dose-fractionated image stacks, which were motion corrected using UCSF MotionCor2[7] and analyzed with RELION v.3.0.8[8]. Motion corrected images without dose weighting were used for contrast transfer function (CTF) estimation using CTFFIND v.4.1[9] and template-based particle picking was done with Gautomatch v.0.53 (http://www.mrc-lmb.cam.ac.uk/kzhang/) with the Hsp90:p23 crystal structure (2CG9) as a reference to select a total of 718,080 particles. Multiple rounds of 3D classification were performed with 2CG9 as a low pass filtered (40 Å) initial model until a medium-resolution (~8Å) GR:Hsp90:p23 reconstruction was obtained from 13,570 particles. This reconstruction was used as a reference for the larger dataset.

The larger dataset consisted of 5,608 dose-fractionated image stacks, which were motion corrected using UCSF MotionCor2 and analyzed with RELION v.3.0.8. Motion corrected images with dose weighting were used for contrast transfer function (CTF) estimation using CTFFIND v.4.1 and reference-free particle picking was done with RELION v.3.0.8 Laplacian-of-Gaussian auto-picking to select a total of 6,062,152 particles. The processing scheme is depicted in **Supplementary Fig. 2a**. An initial round of three-dimensional (3D) classification was performed without symmetry using a reference model from a previously collected smaller dataset (see above). The class with clearly recognizable Hsp90 density was used for a second round of 3D classification. In this second round, one class with just Hsp90 density was refined to a nominal resolution of 2.78Å and one class with Hsp90:p23 density was refined to a nominal resolution of 2.78Å. The classes with clearly recognizable GR density were used for a third round of 3D classification. Particles from classes with the best GR density were then combined (~340,000 particles) and refined. To improve the resolution of GR and the p23 tail helix, these regions were further refined using focused classification with a mask including GR and the p23 tail helix. The best focused classes were combined (140,217 particles) and refined to a nominal resolution of 2.56Å. Using the 2.56 Å reconstruction, per-particle CTF and beam-tilt was refined using RELION. Although the FSC showed slightly improvement over the pre-refined reconstruction at medium resolution range (5–10Å), the nominal resolution at 0.143 FSC remained unchanged. Nevertheless, we used the CTF/beam-tilt refined particles for the following focused refinement on GR:p23 tail helix and for the resulting reconstructions used for model building. To further improve the resolution of GR and the p23 tail helix for model building, these regions were refined using focused refinement with a mask including GR and the p23 tail helix. From the third round of 3D classification, particles from 3D classes with MBP density were combined (~650,000) and refined. To improve the resolution of MBP, the MBP region was further refined using focused classification with a mask on MBP. The best focused 3D classes were combined (31,556 particles) and refined to a nominal resolution of 3.96 Å.

All final reconstructions were post-processed in RELION in which the nominal resolution was determined by the gold standard Fourier shell correlation (FSC) using the 0.143 criterion (**Supplementary Fig. 2b**). Maps were sharpened and filtered automatically determined by RELION according to an estimated overall map B-factor and filtered to their estimated resolution. RELION was used to estimate the local resolution of each map (**Supplementary Fig. 2a**).

*Model building and refinement*

For the GR-maturation complex atomic model, the dexamethasone-bound human GR crystal structure (1M2Z), the human p23 crystal structure (1EJF), and a homolog model of human Hsp90α was derived from the human Hsp90β from the Hsp90:Cdk4:Cdc37 cryo-EM structure (5FWK) with the sequence alignment (86% sequence identity) obtained from HHpred server[10] were used as the starting models for model building (**Supplementary Table 1**). Models were refined using Rosetta throughout. Following the split map approach[11] to prevent and monitor overfitting, the Rosetta iterative backbone rebuilding procedure was used to refine models against one of the half maps obtained from RELION, with the other half map only used for validations. The structurally uncharacterized $p23_{tail\text{-}helix}$ was first *de novo* built into the density using RosettaCM[12] and then was further refined using the same Rosetta iterative backbone rebuilding procedure. With a proper density weight obtained using the half maps, the final model of the GR:Hsp90:p23 complex was refined against the full reconstruction allowing only sidechain and small-scale backbone refinement. The final refinement statistics are provided (**Supplementary Table 1**). For the Hsp90:p23 and MBP:Hsp90:p23 map densities (**Supplementary Fig. 9a** and **Supplementary Fig. 10a**), the Hsp90:p23 atomic model from the GR-maturation complex was docked into the map densities. For MBP:Hsp90:p23, the apo MBP crystal structure[13] (1OMP) was docked into the map density. In **Supplementary Fig. 10c**, the maltose-bound MBP crystal structure[14] (1ANF) is docked into the map density for comparison.

*Fluorescence polarization assays*

Fluorescence polarization of fluorescent dexamethasone (F-dex)(Life Technologies) was measured on a SpectraMax M5 plate reader (Molecular Devices) with excitation/emission wavelengths of 485/538 nm, temperature control set at 25°C. Buffer conditions were 50 mM HEPES pH 8, 100 mM KCl, 2 mM DTT. For equilibrium ligand binding in **Fig. 3e** and **Supplementary Fig. 8b**, proteins were pre-equilibrated together at room temperature for 60 minutes. Proteins and reagents were added at the following concentration: 250 nM GR, 2 μM Hsp40, 15 μM Hsp70, 15 μM Hsp90, 15 μM Hop, 15 μM p23 or p23 tail mutants, and 5 mM ATP/$MgCl_2$. For **Supplementary Fig. 8b**, proteins were added at the following concentration: 250nM GR and 15μM p23 or p23 tail mutants. Ligand binding was initiated with 20nM F-dex and association was measured until reaching saturation. Plotted equilibrium values represent the mean of 3 separate experiments for **Fig. 3e**, with error bars representing the standard deviation, and 5 independent experiments for **Supplementary Fig. 8b**. For **Supplementary Fig. 8b**, polarization values were normalized to the equilibrium ligand binding of 250nM GR and 20nM F-dex. GR ligand binding behavior was affected by buffer conditions; therefore, reactions were always normalized such that each reaction had equivalent amounts of buffer reagents.

*Sequence alignments and $p23_{tail\text{-}helix}$ motif search*

For the p23 sequence alignments in **Fig. 3c** and **Supplementary Fig. 7d**, sequences were aligned in Clustal Omega[15] and visualized in JalView 2.11.1.0[16]. Sequences in the alignment are: *H. sapiens* p23, *M. musculus* p23, *R. norvegicus* p23, *G. gallus* p23, *X. troicalis* p23, *D. melanogaster* p23, *A. thaliana* p23, and *S. cerevisiae* p23 (Uniprot accession numbers: Q15185, Q9R0Q7, P83868, Q90955, Q5U4Z0, Q7SZQ8, A0A0B4K6D2,Q8L7U4, P28707, respectively). For **Fig. 3d**, the ConSurf server[17,18] was used to select and align 87 GR sequences. The human GR crystal structure[19] (4P6X) was used to select sequences from UNIREF90 with maximal percent ID at 95% and minimal percent ID at 65%. Conservation scores were calculated and provided by the server. The conservation scores calculated by ConSurf were mapped onto GR from the maturation complex atomic model using Chimera. For **Supplementary Fig. 7c**, sequences for the human steroid hormone receptors were aligned in Clustal Omega and mapped onto GR from the maturation complex using Chimera. Conservation was calculated using AL2CO[20] parameters (unweighted frequency estimation and entropy-based conservation measurement). Relating to **Fig. 3c** and **Supplementary Fig. 3d**, the $p23_{tail\text{-}helix}$ motif search was performed using ScanProsite[21]. The motif "FXXMMN" was used to search the UniProtKB sequence database with taxonomy restricted to *Homo Sapiens*. There were 10 total hits on the motif, which included p23 and NCoA3/SRC-3.
